# Pro-inflammatory Cytokine GM-CSF Improves Learning/Memory and Brain Pathology in Dp16 Down Syndrome Mice and Improves Learning/Memory in Wild-Type Mice

**DOI:** 10.1101/2021.07.04.451073

**Authors:** Md. Mahiuddin Ahmed, Athena Ching-Jung Wang, Mihret Elos, Heidi J. Chial, Stefan Sillau, D. Adriana Solano, Christina Coughlan, Leila Aghili, Paige Anton, Neil Markham, Vanesa Adame, Katheleen J. Gardiner, Timothy D. Boyd, Huntington Potter

## Abstract

Down syndrome (DS) is characterized by chronic neuroinflammation, peripheral inflammation, astrogliosis, imbalanced excitatory/inhibitory neuronal function, and cognitive deficits in both humans and mouse models. Suppression of inflammation has been proposed as a therapeutic approach to treating DS co-morbidities, including intellectual disability (DS/ID). Conversely, we discovered previously that treatment with the pro-inflammatory cytokine granulocyte-macrophage colony-stimulating factor (GM-CSF) improved cognition and reduced biomarkers of brain pathology in humans with Alzheimer’s disease (AD), another inflammatory disorder, and in a mouse model of AD. To investigate the effects of GM-CSF treatment on DS/ID, we assessed behavior and brain pathology in 12-14 month-old DS mice (Dp[16]1Yey) and their wild-type (WT) littermates, neither of which develop amyloid, and found that GM-CSF treatment improved performance in the radial arm water maze in both Dp16 and WT mice compared to placebo. Dp16 mice also showed abnormal astrocyte morphology and aggregation and fewer calretinin-positive interneurons, both of which were improved by GM-CSF treatment. These findings suggest that stimulating and/or modulating inflammation and the innate immune system with GM-CSF treatment may enhance cognition in both people with DS/ID and in the typical aging population.

## Introduction

Down syndrome (DS) is a chromosomal disorder and the most common genetic cause of both intellectual disability (ID) and age-associated cognitive decline (1–3), with a worldwide incidence of approximately one in 700-1,000 live births (4–6). DS is caused by an extra copy of all or part of human chromosome 21 (Hsa21), which usually arises due to chromosome 21 non-disjunction during female meiosis, resulting in an egg with two copies of chromosome 21 (Hsa21), which, when fertilized, leads to the presence of trisomy 21 in all or most cells during development and after birth (7, 8). Phenotypes associated with DS involve multiple organs and tissues and vary in incidence and severity among individuals (1, 2, 9, 10). In particular, reduced brain size and neurological abnormalities, including impaired neurogenesis and oligodendrogenesis, decreased synaptogenesis, imbalanced excitatory/inhibitory neuronal activity, increased gliogenesis, altered astrocyte morphology and signaling, and astrogliosis are common in people with DS and are directly associated with ID (11–17). In addition, every person with DS will develop Alzheimer’s disease (AD) brain pathology by age 40, and most will exhibit AD-like dementia by age 60 (18–24). Lastly, compared to the typical population, people with DS are at increased risk for other nervous system abnormalities, such as seizures and autism spectrum disorder (20, 25). A combination of improved medical care, the development of targeted public education, and legislative and social initiatives has led to a dramatic increase in the life expectancy for people with DS, from an average of approximately 10 years in the 1960s to an average of approximately 60 years today (26). There is, therefore, a significant need for safe therapeutic treatments to improve the cognitive abilities of people with DS so that they may enjoy expanded educational and employment opportunities, participate more fully in society, and live more independently.

The cellular and molecular bases of the many diverse features of DS in both the brain and periphery are not well understood. One hypothesis that has received increasing attention and is supported by several lines of evidence is that people with DS are in a state of chronic abnormal inflammation, including features of auto-inflammation. For example, analyses of a selected proteome in both plasma and brain tissue samples from people with DS have revealed dysregulation of inflammatory protein expression, such as striking increases in several pro-inflammatory cytokines and decreases in numerous complement cascade components (27–29). Another study revealed elevated levels of not only pro-inflammatory, but also anti-inflammatory, cytokines in plasma from children with DS (30, 31). These results were recently confirmed and expanded in a comprehensive analysis that showed elevated levels of both pro- and anti-inflammatory markers in the frontal cortex and hippocampi of all ages of people with DS, as well as in primary cultures of DS fetal brain tissue (32). Analyses of protein expression in the brains of mouse models of DS and in lymphoblastoid cell lines from people with DS also show upregulation of inflammation-related genes (33–36).

Relevant to such observations of dysregulated immune responses in DS is the fact that four out of the six genes encoding interferon receptors reside on chromosome 21, thus potentially explaining the observation that expression of interferon-sensitive genes, as well as other downstream inflammatory genes, are upregulated in plasma and/or CSF samples from people with DS (28, 37, 38). The microRNA miR-155, which has been shown to promote inflammatory responses (39), also maps to Hsa21 and is overexpressed in people with DS.

Together, these findings have led investigators to attribute many features of people with DS to their state of chronic abnormal inflammation and to propose that suppressing inflammation may have therapeutic benefits across many features of the disorder (24, 27–29, 32, 40).

Observations of elevated levels of both pro- and anti-inflammatory biomarkers in DS appear contradictory and require careful consideration of which abnormalities are detrimental and which may be compensatory. For example, in contrast to the view that inflammation in DS is a key target for therapeutic intervention, we propose and have tested an alternative hypothesis that recruiting the innate immune system and its inflammatory features may successfully ameliorate at least the cognitive deficits in DS. This hypothesis is based in part on previous work in which we showed that: (1) treatment with the inflammatory cytokine and innate immune system modulator, granulocyte-macrophage colony-stimulating factor (GM-CSF), improves cognition in both a mouse model of AD and in middle-aged wild-type (WT) mice (41), observations that were later replicated by other groups (42, 43), (2) treatment with recombinant human GM-CSF (sargramostim/Leukine^®^) was associated with improved cognition in leukemia patients with chemotherapy-associated cognitive impairment after chemo-ablation and hematopoietic stem cell transplantation (44), and (3) treatment with GM-CSF/sargramostim was safe, significantly improved Mini-Mental State Exam (MMSE) scores, and partly restored levels of plasma biomarkers of AD brain pathology and neuronal damage (Aβ40 and Aβ42, total tau, and ubiquitin C-terminal hydrolase L1 [UCH-L1]) toward normal in participants with mild-to-moderate AD after only 15 days of subcutaneous injections in our recently completed Phase 2 clinical trial [NCT01409915, the results of which have been published (45). Based on these observations, we hypothesized that GM-CSF may act as a general cognition enhancer, despite and possibly because of its pro-inflammatory activity, which could potentially ameliorate the cognitive deficits observed in mouse models of DS, with potential implications for treating people of all ages with DS. In the current study, we tested our novel hypothesis, which is in contrast to the commonly held view that inflammation contributes to DS-related, and possibly age-associated, deficits in cognition and therefore should be suppressed to effect therapy.

The long arm of Hsa21 harbors ∼165 traditional protein-coding genes, ∼50 genes encoding members of the family of keratin-associated proteins, a small number of characterized functional RNA genes, and several hundred genes that may be functional RNAs, protein-coding, or transcriptional noise (46). Although the functional annotation of these genes is far from extensive, several have been shown to play roles in brain development and/or to lead to learning/memory deficits when their gene expression levels are altered (36). When present in three copies or when mutated, the *APP* amyloid precursor protein gene, which resides on Hsa21, causes familial AD (FAD). APP is likely to play a key role in the development of AD in DS, although other Hsa21-encoded proteins may also contribute (22, 24, 36).

Animal models are essential for developing and testing therapeutic interventions for DS. It has proven challenging to generate a mouse model that is trisomic for orthologs of all Hsa21 genes because the orthologs map to regions of three mouse chromosomes: Mmu16, Mmu17, and Mmu10. The Mmu16 segment has received the most attention in DS mouse models, in part because of the large number of Hsa21 protein-coding orthologs (i.e., 102) that it harbors. The first viable, and currently the most well-studied, segmental trisomy mouse model of DS, the Ts65Dn mouse, is trisomic for 90 of the 102 Mmu16 orthologs. It also exhibits a number of phenotypes relevant to DS, including learning/memory deficits, thus emphasizing the contributions of Mmu16 orthologs to ID in DS. However, the Ts65Dn mouse is also trisomic for ∼35 protein-coding genes whose orthologs do not reside on Hsa21 and are not trisomic in people with DS; thus, the presence of those extra genes may confound the interpretation of Ts65Dn phenotypes (46). Therefore, to test our hypothesis that GM-CSF may be used to treat cognitive deficits in people with DS, we employed the Dp(16)1Yey/+ mouse model of DS (referred to herein as Dp16), which is trisomic for ∼102 orthologs of Hsa21 protein-coding genes, including the entirety of the Mmu16 orthologous segment, and it is not trisomic for any other genes (47). Similar to other mouse models of DS, Dp16 mice do not develop AD-related neuropathology, such as amyloid plaques, despite harboring three copies of the mouse *App* gene, because the mouse Aβpeptide exhibits little propensity to aggregate into neurotoxic β-sheet oligomers and fibrils (48). Thus, Dp16 mice serve as a model of DS, but not of AD, and are therefore ideal for pre-clinical testing of our amyloid-independent alternative hypothesis that treatment with GM-CSF/sargramostim will improve cognition in people with DS. In the current study, we treated Dp16 mice and their WT littermates with GM-CSF or with placebo, assessed their cell and biomarker innate immune system/inflammatory response, their learning/memory performance, and then compared brain pathology in astrocytes and interneurons by indirect immunofluorescence microscopy staining.

## Methods and Materials

### Animals

The Dp(16)1Yey (abbreviated Dp16) mouse model of DS was generated using Cre/loxP-mediated chromosomal rearrangement to duplicate a 22.9 Mb segment of Mmu16 spanning the segment syntenic with Hsa21 (47). Male Dp16 mice on the C57BL/6J background were purchased from Jackson Laboratory (Stock # 013530) and crossed with female C57BL/6J wild type (WT) mice to generate the litters used for these experiments. All animals were bred at the University of Colorado Anschutz Medical Campus and maintained in a room with HEPA-filtered air and a 14:10 light:dark cycle, fed a 6% fat diet, and provided with acidified (pH 2.5-3.0) water *ad libitum*. All procedures were approved by the University of Colorado Institutional Animal Care and Use Committee (IACUC) and were performed in accordance with National Institutes of Health guidelines for the care and use of animals in research. Same-sex littermates were housed in the same cage after weaning. Mice were genotyped by polymerase chain reaction (PCR) using a protocol from the Jackson Laboratory (Bar Harbor, ME). For the behavioral and brain pathology experiments, we started with 19 WT male mice, 11 Dp16 male mice, 13 WT female mice, and 12 Dp16 female mice, but two of the WT male mice did not perform the RAWM task at the baseline assessment (they floated), and only the remaining 17 WT male mice were included in the post-treatment analyses. All physically healthy-looking WT mice were chosen from their littermates. All behavioral and brain pathology experiments were conducted with mice aged 12-14 months. Open Field and Y-maze experiments used both male and female Dp16 mice and their age- and sex-matched WT littermate controls; all other behavioral and brain pathology experiments used only male mice. Open Field and Y-maze tasks were performed between 9:00 AM and 11:00 AM, and the RAWM maze was assessed from 9:00 AM to 1:00 PM because the RAWM requires more time to complete the task. For studies of plasma biomarkers, we used another set of mice, including 7 WT male mice, 8 Dp16 male mice, 3 WT female mice, and 2 Dp16 female mice, aged 12-15 months.

### Drug Administration

All of the mice in the behavioral and brain pathology studies were injected subcutaneously five days/week, Monday-Friday, for five weeks with GM-CSF (5 µg/day) or with an equal volume of saline (200 μl/day) for a total of 24 injections over a period of 32 days including non-treatment days. As discussed, two WT male mice did not perform the RAWM task at the baseline assessment (they merely floated in place), and those two mice were not included in the post-treatment analyses. Mice were divided into four treatment groups: (1) WT mice injected with saline, including 9 males and 6 females; (2) WT mice injected with GM-CSF, including 8 males and 7 females; (3) Dp16 mice injected with saline, including 5 males and 6 females; and (4) Dp16 mice injected with GM-CSF, including 6 males and 6 females (Figure 1 shows the study timeline and assessments). For studies of plasma biomarkers, we used another set of mice, including 7 WT male mice, 8 Dp16 male mice, 3 WT female mice, and 2 Dp16 female mice, aged 12-15 months, that were injected with GM-CSF (5 µg/day) or with an equal volume of saline (200 μl/day) for 10 consecutive days (see Measurements of Plasma Biomarkers below).

**Figure 1.**
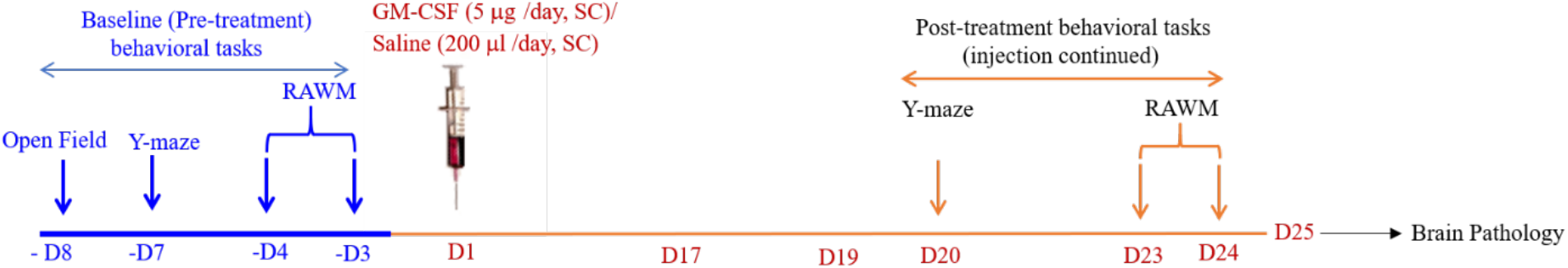
Schematic of the Study Timeline. Each day (D1-D25) refers to a treatment day. After the baseline pre-treatment assessments, the cohorts of Dp16 and WT control mice were injected five days/week with either GM-CSF (5 μg/day) or an equal volume of saline (200 μl/day), Monday-Friday, for five weeks for a total of 24 injections (ending on D24) over a period of 32 days including non-treatment days. There were four treatment groups for both female and male mice: (1) WT mice injected with saline (9 males; 6 females), (2) WT mice injected with GM-CSF (8 males; 7 females), (3) Dp16 mice injected with saline (5 males; 6 females), and (4) Dp16 mice injected with GM-CSF (6 males; 6 females). Both male and female mice were tested in the Open Field task at baseline and in the Y-maze task at baseline and post-treatment. Only male mice were used for the RAWM task at baseline and post-treatment because the female mice had poor swimming abilities at this age.

### Behavioral Assessments

For all behavioral/activity assessments, on the day of testing, the mice were brought into the testing room and allowed to habituate for 30 min in their home cage.

### Open Field

To evaluate locomotor and exploratory behavior, in naïve mice, each animal was placed in the center of an arena (44 cm x 44 cm) and allowed to explore for 10 min with the room lights on. Activity was recorded by a video tracking system using EthoVision XT software (Noldus). Total distance traveled, total time spent in the center zone, frequency of presence in the center zone, and speed of movement were all analyzed during the 10 min session.

### Spontaneous Alternation Y-Maze

The spontaneous alternation Y-maze assesses spatial working memory. The apparatus is composed of three arms set at 120-degree angles from a center area. The arms are 40 cm long, 8 cm wide, and 15 cm high. The mouse was placed in the center area and allowed to explore freely; the session was stopped after the mouse had completed 20 arm entries or had explored for 8 min (if the mouse did not complete 20 entries within 8 min, we counted the total number of entries in 8 min). The EthoVision XT software and video tracking system was used to record the number of arms visited. Spatial working memory was measured as the percentage of the number of successful alternations (sequential entries into all three arms) divided by the total number of arm entries.

### Baseline Radial Arm Water Maze (RAWM) Assessments

The RAWM is a hippocampal-based task measuring working memory. Only male mice were used for the RAWM because 12-14 month-old female mice had poor swimming abilities. The RAWM (49) was first performed as a baseline assessment at six days before the first injection of either GM-CSF or saline. The RAWM apparatus consists of six arms and is placed in a circular pool of water. A two-day RAWM task was performed in which a submerged (hidden) platform was placed in arm 4 throughout testing. The maze is located in a room with numerous cues on the walls and surroundings that the mice can use to orient within the maze.

On both Day 1 (learning/training) and Day 2 (testing), each mouse was exposed to 15 trials (60 sec each), run in three sessions: 1^st^ session: 6 trials with 5 min rest between trials; 50 min rest before session 2; 2^nd^ session: 6 trials with 5 min rest between trials; 50 min rest before session 3; and 3^rd^ session: 3 trials with 5 min break between trials. Before the first trial, each mouse was allowed to spend 30 sec on the platform to habituate to the location of the platform with respect to the walls and room surroundings. During each trial, the mouse was started from a different arm using a randomization scheme such that during the 15 trials, the mouse started in each of the five arms without the platform three times. On Day 1 (training day), each mouse was gently placed into one of the five arms (excluding Arm 4, which contained the platform) and was allowed a maximum of 60 sec to find the platform; if the platform was not found during this time, the mouse was gently guided to it. The mouse remained on the platform for 10 sec and was then gently towel-dried and transferred to the home cage located on a heating pad, for a 5 min break between trials. Between sessions, the mouse rested for 50 min in the home cage.

On Day 2 (testing day), spatial memory was tested with the platform still located in Arm 4. Similar to Day 1, each mouse was gently placed into one of the five vacant arms and again allowed a maximum of 60 sec to find the platform. As described for Day 1, 15 trials were performed with each mouse, run as three sessions with a 50 min break between sessions. Mouse behavior was recorded with a video camera located directly above the water pool. Both the number of errors (incorrect arm choices) and the escape latency were recorded for each trial. For analysis purposes, both Day 1 and Day 2 data from the RAWM were used separately in baseline assessments. After averaging the data from 15 trials using the time values for each individual mouse, the mean time to reach the platform between Dp16 and WT control mice was compared on both days.

### Post-Treatment Alternating Y-maze

After the baseline pre-treatment behavioral testing, the cohorts of Dp16 and WT littermates were injected with either GM-CSF or saline. Post-treatment Y-maze performances of male and female mice were measured starting on treatment day 20 using the same protocol as for baseline measurements.

### Post-Treatment RAWM Tasks

Male mice were evaluated post-treatment in the RAWM starting on treatment day 21. The protocol was the same as for baseline measurements, with the exception that on Day 2, the platform was located in Arm 6, instead of in Arm 4. To evaluate memory function post-treatment, the latency of each saline- or GM-CSF-injected mouse on post-treatment day 1 was normalized with its mean latency at block 5 of pre-treatment day 2 (baseline day 2), and the data were calculated relative to 100% of block 5 pre-treatment day 2. Normalized latencies were analyzed in blocks of three trials (for a total of five blocks from 15 trials), and the GM-CSF-injected group was compared with the saline-injected group. For quantitative analyses, averaging the normalized latency of post-treatment day 1 from blocks 1-5 and combining blocks 1-2 or blocks 4-5 were compared between GM-CSF- and saline-injected mice for each genotype. To assess the effects of GM-CSF treatment over time, the combined normalized latency of blocks 1-2 were compared with the combined normalized latency of blocks 4-5 in GM-CSF-treated Dp16 mice.

On Day 2 of the post-treatment RAWM testing, the platform was moved to a new arm (from Arm 4 to Arm 6) while other maze cues remained unchanged, and then each mouse performed 15 trials (60 seconds each) to evaluate learning flexibility. To assess learning flexibility, the latency of each saline- or GM-CSF-injected mouse at post-treatment day 2 was normalized with its mean latency at block 5 of post-treatment day 1, and the data were calculated relative to 100% of block 5 post-treatment day 1. Normalized latencies from post-treatment day 2 were analyzed in blocks of three trials (for a total of five blocks from 15 trials), and the GM-CSF-injected group was compared with the saline-injected group. For quantitative analyses, averaging the normalized latencies of post-treatment day 2 from all blocks 1-5 were compared between GM-CSF- and saline-injected mice for each genotype.

### Measurements of Plasma Biomarkers

For studies of plasma biomarkers, we used another set of mice (15 males and 5 females, aged 12-15 months) that were injected subcutaneously with GM-CSF (5 µg/day) or with an equal volume of saline (200 μl/day) for 10 consecutive days, with the last injection at 8:00 AM on day 10. The mice were divided into four treatment groups: (1) WT mice injected with saline, including 3 males and 2 females; (2) WT mice injected with GM-CSF, including 4 males and 1 female; (3) Dp16 mice injected with saline, including 4 males and 1 female; and (4) Dp16 mice injected with GM-CSF, including 4 males and 1 female. Blood samples were collected at 4-6 h after the last injection on day 10 in microtubes with EDTA (Sarstedt Inc., Nümbrecht, Germany) from cardiac puncture under anesthesia. The blood samples were centrifuged at 1,500 RCF for 15 min at room temperature to separate the plasma. The plasma was then transferred to a clean microtube and centrifuged a second time at 2,200 RCF for 10 min at 4°C. The resultant plasma was collected and stored at −80°C until use.

We used a custom U-PLEX Metabolic Group 1 (Mouse) Multiplex Assay (Meso Scale Discovery, Rockville, MD, catalog number K152ACL-1) designed to measure 23 different analytes, including cytokines and metabolic proteins, in mouse plasma samples. Results from eight of the analytes were below the limits of detection, but we were able to measure 15 analytes, including IL-6, IL-2, TNF*α*, IP-10, MCP-1, MDC, IL-1β, IL-5, IL-22, IL-33, IFN*γ*, VEGF-A, KC/GRO, BDNF, and MMP-9. We used the U-PLEX plate to measure the plasma levels of the analytes according to the manufacturer’s instructions. Briefly, biotinylated capture antibodies were mixed with specific U-PLEX Linkers, added to the appropriate wells, and incubated at room temperature for 30 min. The U-PLEX plates were then washed using an automated plate washer (BioTek ELX5012). Plasma was defrosted, diluted two-fold (Diluent 13 buffer, Meso Scale Discovery), and centrifuged at 2,000 x g for 3 min. Fifty microliters of calibrators or plasma were then added to the wells in triplicate, and the plates were incubated for 2 h at room temperature on a Compact Digital Microplate Shaker (ThermoFisher) at 600 rpm. The plates were washed, and 50 μl of diluted detection antibodies were added and incubated for 1 h at room temperature. After washing, MSD GOLD Read Buffer B (Meso Scale Discovery) was added, the plates were read immediately on a MESO QuickPlex SQ 120 instrument (Meso Scale Discovery), and the data were analyzed using DISCOVERY WORKBENCH software (Meso Scale Discovery).

### Tissue Processing

Following the post-treatment assessments, on post-treatment day 25, the mice that were assessed in the RAWM (males only) were anesthetized with sodium pentobarbital, perfused intracardially with saline for 5 min, and the brains were then removed rapidly. The left hemisphere was quickly dissected into hippocampus, cerebellum, frontal cortex, and remaining cortex (i.e., total cortex minus frontal cortex), frozen immediately in liquid nitrogen, and then stored at −80°C until use. The right hemisphere was immersed in freshly prepared 4% paraformaldehyde (PFA) in PBS for 24 h at 4°C. After fixing with PFA, 4 μm-thick paraffin-embedded hippocampal brain sections were mounted on glass slides and used for indirect immunofluorescence microscopy staining or stored at 4°C until use.

### Immunohistochemistry and Image Analyses

Immunofluorescence microscopy analyses were carried out using 4 μm-thick, paraffin-embedded hippocampal brain sections from male Dp16 and WT mice treated with GM-CSF or saline. Briefly, after standard deparaffination, the slides were incubated for 30 min with just-boiled 10 mM sodium citrate buffer for antigen retrieval. The slides were washed three times (3X) in TBST and incubated in blocking solution (Protein Block Serum-Free, Agilent DAKO, CA, USA) for 30 min at room temperature (RT), followed by incubation with rabbit anti-GFAP primary antibody at a 1:200 dilution (Abcam, MA, USA) in blocking solution, or with a rabbit anti-calretinin primary antibody at dilution of 1:200 (Swant, Switzerland) in blocking solution overnight at RT. The following day, the slides were washed 3X in TBST with gentle rocking and incubated with Alexa-Flour488-conjugated anti-rabbit secondary antibody (Invitrogen, CA, USA). The slides were mounted using 4,6-diamidino-2-phenylindole (DAPI) Fluoromount-G (SouthernBiotech) and sealed with a coverslip. The preparation and staining of sections from each mouse brain were repeated with two slides, which yielded similar results. Each slide contained three brain sections from each mouse; two sections were incubated with a specific primary antibody, and one section was incubated in blocking solution without antibody as a negative control. There were four immunostained brain sections from each mouse. All sections were imaged using an Olympus IX83 Fluorescence microscope (Center Valley, PA). All images were taken at 20X magnification using the same exposure time. For quantitative analyses, the number of GFAP-positive astrocyte clusters and calretinin-positive interneurons were counted manually within brain regions of interest using cellSens software from Olympus by two different experimenters who were blinded as to genotype and treatment. Before counting, the quality of all images was examined carefully, and only images without broken/folded or smeared tissues were used. At least two sections from each mouse (at least one section from each slide) were usable, for a total of 2-4 brain sections per mouse. After averaging the number of GFAP-positive astrocyte clusters for each mouse, the mean number of clusters was compared between Dp16 and WT mice treated with saline or GM-CSF in the entire hippocampal region. Similarly, after averaging the number of calretinin-positive interneurons from 2-4 repeats for each mouse, the mean numbers of calretinin-positive interneurons were compared between Dp16 and WT mice treated with saline or GM-CSF in the entorhinal cortex and subiculum.

### Statistical Analyses

Statistical significance between groups was determined by unpaired Student’s t-test and by ANOVA, followed by Tukey’s multiple comparisons test using GraphPad Prism 7.00. All data were expressed as the mean +/-SEM. A p-value of less than 0.05 was considered statistically significant.

## Results

### Overview and Study Timeline

For baseline measurements, we first evaluated separate cohorts of untreated male and female Dp16 mice and their WT littermates at 12-14 months of age in the Open Field and the Y-maze tasks, and untreated male mice in the radial arm water maze (RAWM), because female Dp16 mice at this age are poor swimmers. We then investigated the effects of GM-CSF treatment versus placebo (saline) on Y-maze and RAWM performance, and on brain pathology. A timeline showing the experimental design is provided (Figure 1).

### Dp16 Male and Female Mice Show Elevated Locomotor Activity in the Open Field Task

Dp16 mice and their WT littermates were first evaluated for genotypic effects on exploratory behavior in the Open Field task. The mice were allowed to explore the Open Field arena for 10 min as a baseline assessment (50). The total distance traveled, total time spent in the center zone, and speed were measured. Both male and female Dp16 mice showed significantly increased distance traveled compared to their WT littermates (*p*=0.026 for male and *p*=0.014 for female genotype comparisons) (Figures 2A and 2D). The Dp16 mice also showed a significantly higher speed compared to their WT littermates (*p*=0.027 for male and *p*=0.014 for female genotype comparisons) (Figures 2C and 2F). However, the average amount of time spent in the center zone was not significantly different between Dp16 mice and their WT littermates, showing that there are no differences in anxiety behavior between the two groups (Figures 2B and 2E). These results show that both male and female Dp16 mice show hyperactivity, but not increased anxiety, in a novel environment.

**Figure 2.**
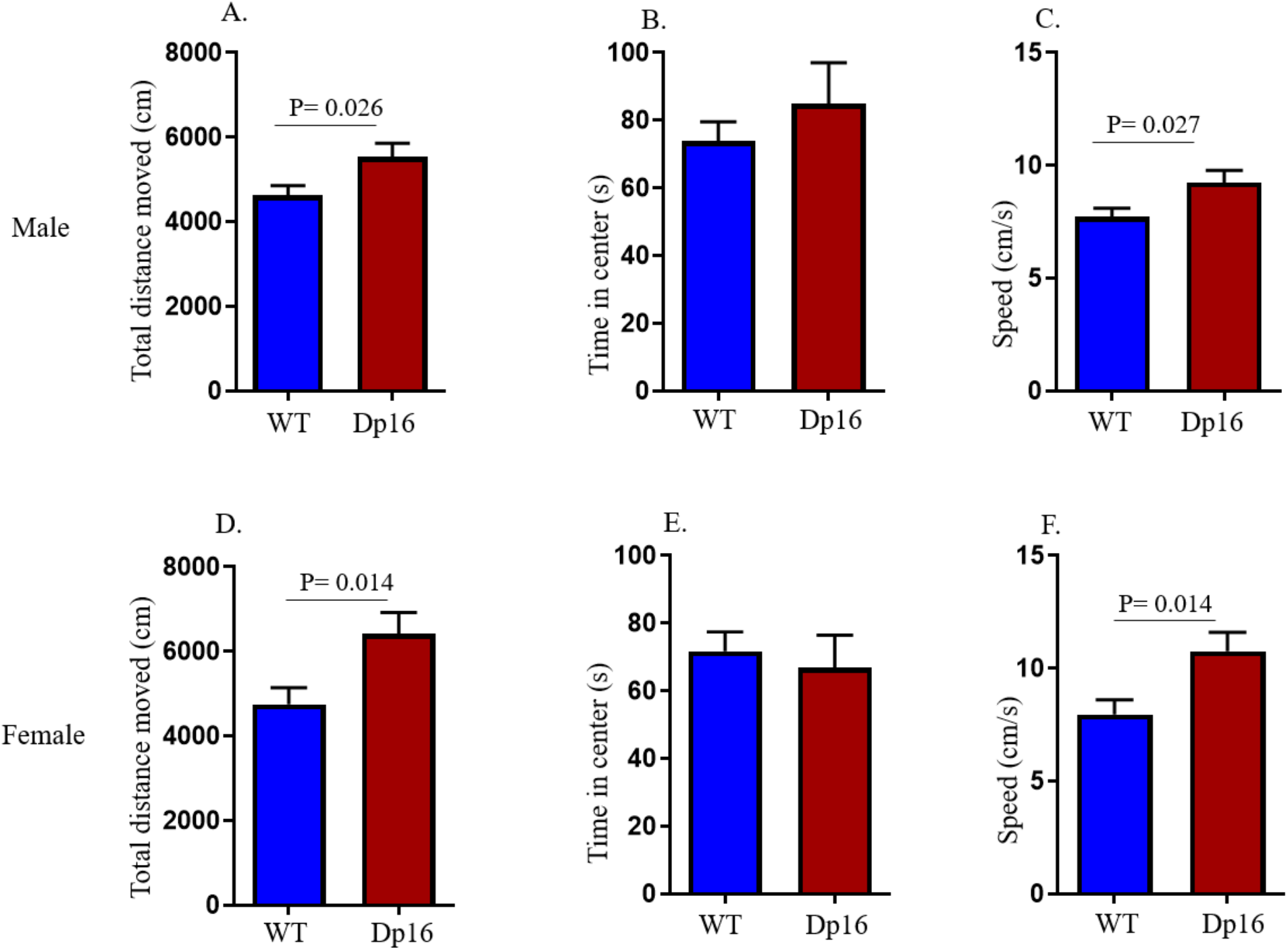
Male and Female Dp16 Mice Show Elevated Locomotor Activity in the Open Field Task at Baseline. Locomotor and exploratory activity were measured in the Open Field task at baseline. Both male and female Dp16 mice showed a significant increase in total distance traveled (A and D, respectively) and speed of movement (C and F, respectively) compared to male and female WT mice, respectively. No significant differences were observed in the time spent in the center for male and female Dp16 mice (B and E, respectively) compared to male and female WT littermates, respectively. Data are represented as mean ± SEM for separate groups of mice (male WT: n=19; female WT: n=13; male Dp16: n=11; and female Dp16: n=13). Statistical significance was determined by the unpaired Student’s *t*-test for comparison between Dp16 and WT littermates.

### Female, but Not Male, Dp16 Mice Show Deficits in the Y-maze Task for Spatial Working Memory at Baseline, and Saline-Injected Male Dp16 Mice Perform Better than GM-CSF-Treated Male Dp16 Mice

On the day following the Open Field test, mice were assessed for spatial working memory in the spontaneous alternation version of the Y-maze, where sequential entry of different arms of the maze assumes better working memory. At baseline, female Dp16 mice showed a significantly lower percentage of alternation compared to their female WT littermates (*p*=0.009) (Figure 3A). Indeed, the 50% alternation rate in female Dp16 mice corresponds to that expected by chance, showing that female Dp16 mice are impaired in working memory. In contrast, Y-maze performance of male Dp16 mice did not differ significantly from that of their male WT littermates (Figure 3B).

**Figure 3.**
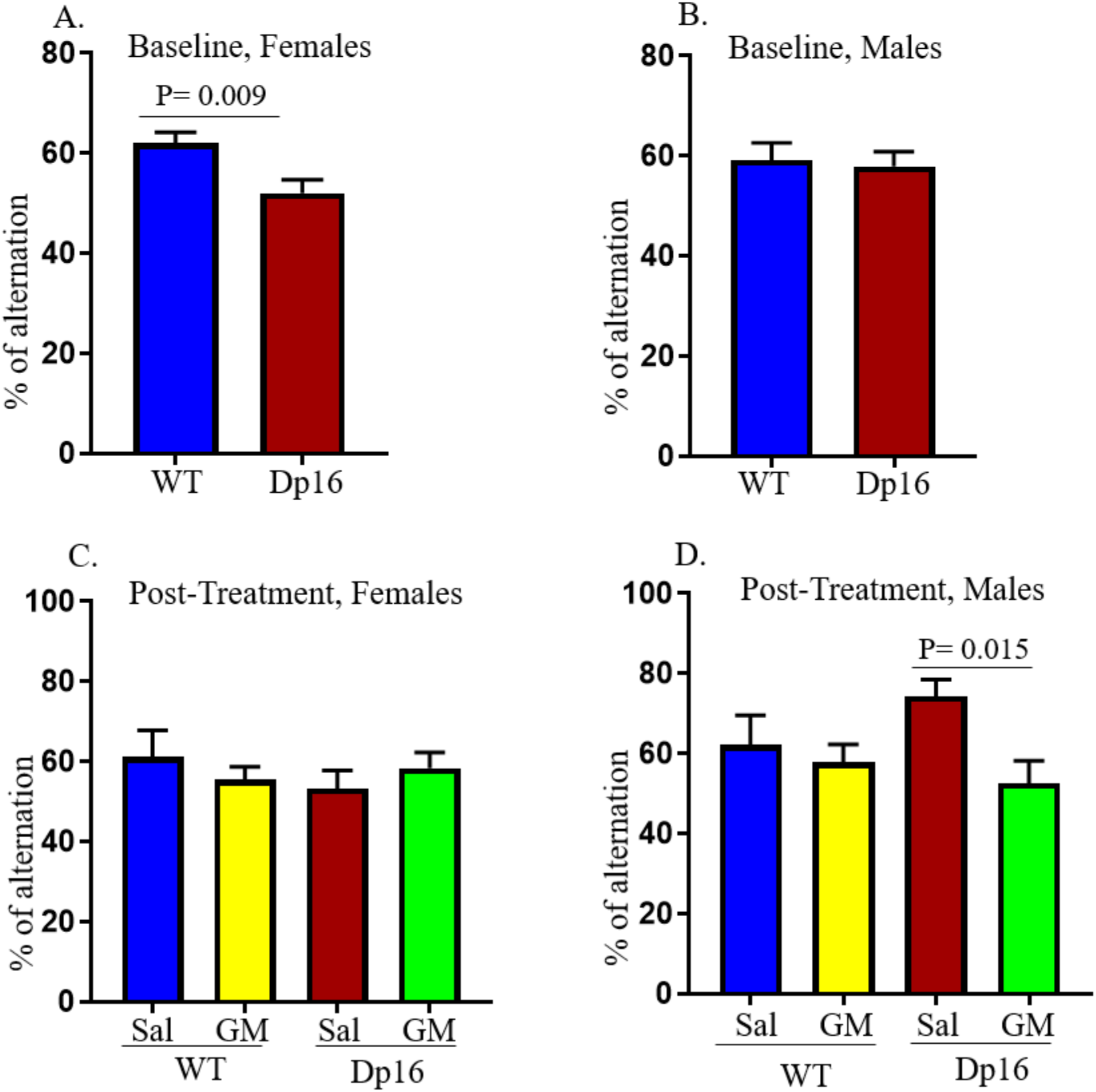
Spontaneous Alternation Rates in the Y-maze Task at Baseline and After GM-CSF Treatment. Female Dp16 mice showed a significantly lower alternation rate in the Y-maze compared to female WT littermates at baseline (A). No significant differences in alternation rates in the Y-maze were observed in male Dp16 mice compared to male WT littermates at baseline (B). No significant differences in spontaneous alternation rates in the Y-maze were observed in GM-CSF-treated female Dp16 mice compared to saline-treated female Dp16 mice (C). GM-CSF-treated (GM) male Dp16 mice show a significantly lower alternation rate in the Y-maze compared to saline-treated (Sal) male Dp16 mice, largely because the saline-treated male Dp16 mice exhibited higher % alternations from baseline levels (D). Data are represented as mean ± SEM for the separate groups of mice, including WT-Sal (9 males; 6 females), WT-GM (8 males; 7 females), Dp16-Sal (5 males; 6 females), and Dp16-GM (6 males; 6 females). Statistical significance was determined by the unpaired Student’s *t*-test for comparison between groups.

Next, the same mice were divided into four treatment groups: (1) WT mice injected with saline, (2) WT mice injected with GM-CSF, (3) Dp16 mice injected with saline, and (4) Dp16 mice injected with GM-CSF (see Figure 1). No significant differences in spontaneous alternation were observed in WT or Dp16 female mice treated with GM-CSF compared to WT or Dp16 female mice injected with saline (Figure 3C). In male mice injected with saline, the performance of Dp16 mice did not differ significantly from that of their WT littermates (Figure 3D). However, Dp16 male mice treated with GM-CSF showed a significantly lower percentage of spontaneous alternations compared to Dp16 male mice injected with saline, largely because the saline-treated Dp16 male mice exhibited an unexpectedly high percentage of alternations compared to baseline (Figures 3B and 3D). Taken together, these findings show that GM-CSF treatment does not rescue working memory deficits in female Dp16 mice, and that GM-CSF treatment does not improve working memory performance in male Dp16 mice as measured by the Y-maze task (Figures 3C and 3D).

### Male Dp16 Mice Show Significant Deficits in Spatial Learning and Memory in the Radial Arm Water Maze (RAWM) Task at Baseline

To evaluate hippocampal-dependent spatial learning and memory, mice were assessed in a two-day RAWM protocol prior to any treatment. Only male mice could be used in these experiments because female Dp16 mice at this age had inadequate swimming abilities. For the baseline assessments, on Day 1, Dp16 mice and their WT littermates were subjected to 15 trials of 60 sec each, and latency to reach the submerged platform was measured. Data were analyzed in blocks of three trials (for a total of five blocks). Latency for both groups decreased over the 15 trials, indicating that they had learned the platform location (Figure 4A). On day two, which served as the memory testing day, both Dp16 and WT mice were again subjected to 15 trials (60 sec each), and the data were analyzed in blocks of three trials. Although Dp16 and WT mice showed no significant differences in memory/latency at the end (block 5) of day 1, they did show a statistically significant difference in the first block of day 2 (p<0.05) (Figures 4A and 4B). The two genotypes showed significant differences in latency between the first four blocks, but they became indistinguishable by block five (Figure 4B).

**Figure 4.**
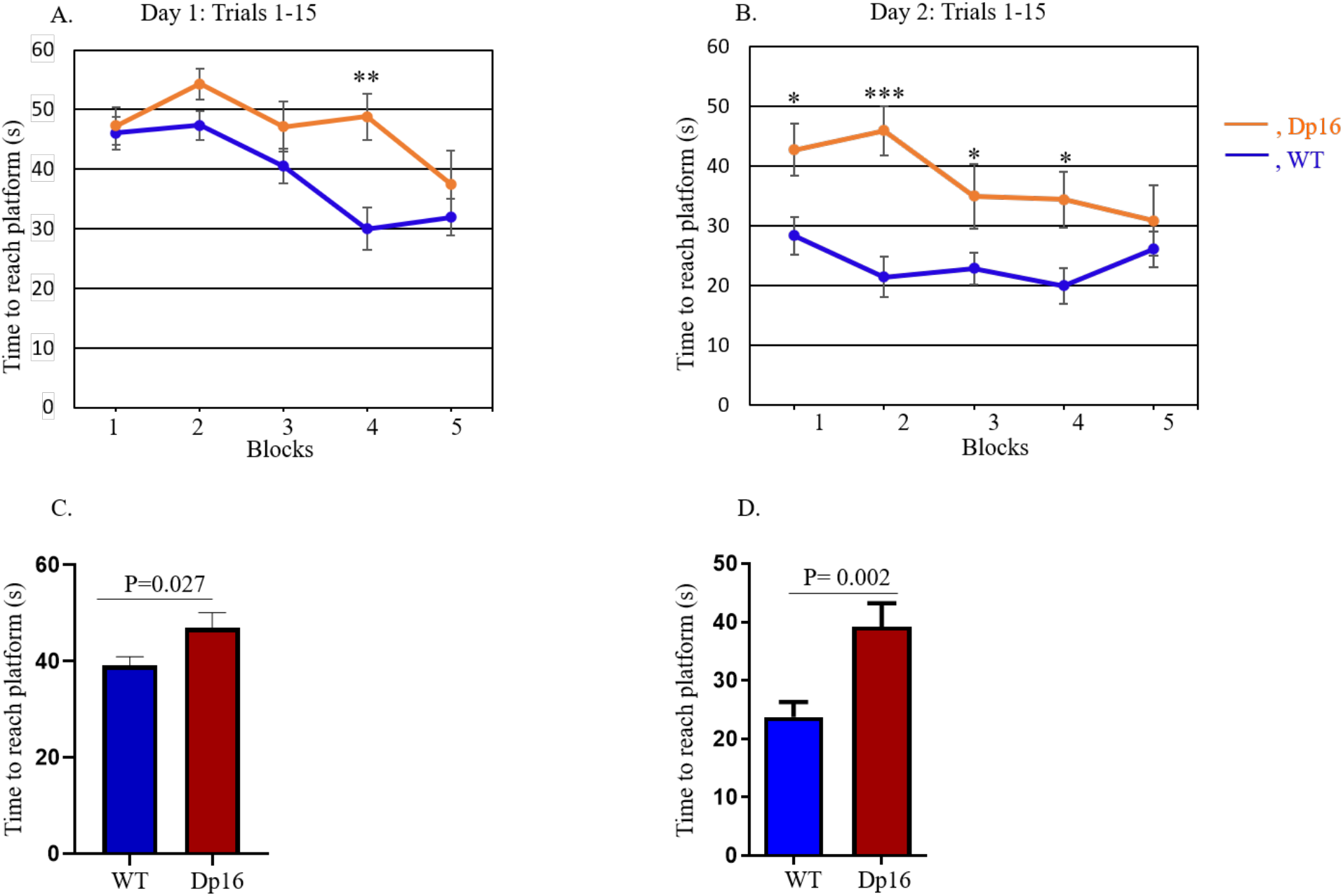
Male Dp16 Mice Show Deficits in the RAWM Task at Baseline. Hippocampal-based spatial learning and memory performance were evaluated in male Dp16 mice for two days using the radial arm water maze (RAWM) task. Data were analyzed considering each group of three out of 15 trials as a block for Day 1, the training day (A), and for Day 2, the testing day (B). On day 2, in blocks 1 and 2, there was a significant difference in memory performance between the Dp16 and WT littermates. After averaging the data from 15 trials using the time values for each individual mouse, the mean time to reach the platform for each day was compared between Dp16 and WT mice for Day 1 (C) and Day 2 (D). Dp16 mice required a significantly longer time to reach the platform than WT mice on both days, although they ultimately learned as well as WT mice, as indicated by non-significant differences in latencies in Block 5. Data are represented as mean ± SEM for separate groups of mice (WT, n=17; Dp16, n=11). Statistical significance was determined by the unpaired Student’s *t*-test for comparison between Dp16 and WT mice (C, D) and by ANOVA followed by Tukey’s multiple comparisons test (A, B). \**p* <0.05; \*\*\**p* <0.001.

Finally, after averaging the data from 15 trials using the time values for each individual mouse, we calculated the mean latency time to reach the platform and compared Dp16 to WT mice on both days. On Day 1, Dp16 mice required a significantly longer time to reach the platform compared to WT controls (*p*=0.027) (Figures 4C). Similarly, on Day 2, Dp16 mice also required a significantly longer time to reach the platform compared to WT controls (*p*=0.002) (Figures 4D). These results show that male Dp16 mice exhibit significant cognitive (both learning and memory) deficits overall in the RAWM task. However, by block 5 of both day 1 and day 2, the latencies of the Dp16 and WT mice were not statistically significantly different. Together, these results show that the main learning deficit in the Dp16 mice was in the speed with which they learned the task. With enough practice, the Dp16 mice eventually performed as well as their WT littermates.

### GM-CSF Treatment Improves Spatial Memory in Male Dp16 and WT Mice in the RAWM Task

We repeated the RAWM analysis in male Dp16 mice and their male WT littermates after treatment with GM-CSF or saline. The male mice were divided into four treatment groups: (1) WT mice injected with saline, (2) WT mice injected with GM-CSF, (3) Dp16 mice injected with saline, and (4) Dp16 mice injected with GM-CSF (Figure 1). We used the same RAWM protocol for baseline assessments, i.e., each mouse performed 15 trials (60 sec/trial) with the hidden/submerged platform placed in Arm 4. Latency for block 5 of pre-treatment day 2 represents maximal learning for baseline assessments; therefore, these values were used for the normalization of post-treatment day 1 latencies. Specifically, we normalized the latencies for each saline- or GM-CSF-injected mouse for each block of post-treatment day 1 with the corresponding latencies from block 5 of pre-treatment day 2 (baseline day 2) and present the values as a percentage of block 5 pre-treatment day 2. Normalized latencies were analyzed in blocks of three trials (for a total of five blocks from 15 trials), and the GM-CSF-treated group was compared with the saline-injected group. Post-treatment latencies in the Dp16 mice treated with GM-CSF showed significant differences from saline-injected Dp16 mice in blocks four and five (Figure 5A). Averaging the normalized latency of post-treatment day 1 from all five blocks (from Figure 5A) showed that the Dp16 mice treated with GM-CSF required significantly less time (shorter latency) to reach the platform compared to the saline-injected Dp16 mice (*p*=0.009) (Figure 5B). To observe the final learning/memory differences over the course of the day in the Dp16 mice injected with GM-CSF or saline, we combined normalized latency of blocks 4 and 5 and compared the GM-CSF-treated groups with the saline-injected groups. Dp16 mice treated with GM-CSF showed significantly shorter latency compared to the saline-injected Dp16 mice (*p*=0.008) (Figure 5C). Furthermore, to clarify the longitudinal learning performance of Dp16 mice treated with GM-CSF, we combined data from blocks 1 and 2 and compared with the combined data from blocks 4 and 5 of Dp16 mice treated with GM-CSF. Figure 5D shows that combined data from the normalized latency of blocks 4 and 5 show significantly smaller latency than the combined latency from blocks 1 and 2 (*p*=0.03), showing that Dp16 mice treated with GM-CSF are improving their learning and memory over time. WT mice treated with GM-CSF required significantly less time to reach the platform in block 1 compared to saline-injected WT mice (Figure 5E). Because learning is a continuous process and both GM-CSF-treated and saline-injected WT mice continued to learn during the day, there are no significant differences in blocks 2-5, indicating that GM-CSF had little effect on performance. Combining the normalized latency of blocks 1 and 2, and then comparing GM-CSF-treated WT mice with the saline-injected WT mice revealed that the WT mice treated with GM-CSF showed significantly smaller latency compared to the saline-injected WT mice for the first two blocks of the day (*p*=0.02) (Figure 5F). Furthermore, even at the very beginning of the task on post-treatment day 1, WT mice treated with GM-CSF showed a significant difference in performance compared to the saline-injected WT mice (Figure 5E, block 1), which remained constant over time, whereas saline-injected WT mice improved in the task and performed similarly to GM-CSF-treated WT mice by the end. To further investigate the learning performance of WT mice injected with saline, we combined data from blocks 1 and 2 and compared them with the combined data from blocks 4 and 5. Figure 5G shows a significantly shorter latency for the results of blocks 4 and 5 compared to that from blocks 1 and 2 (*p*=0.04), showing that WT mice injected with saline improve their learning and memory performance over time.

**Figure 5.**
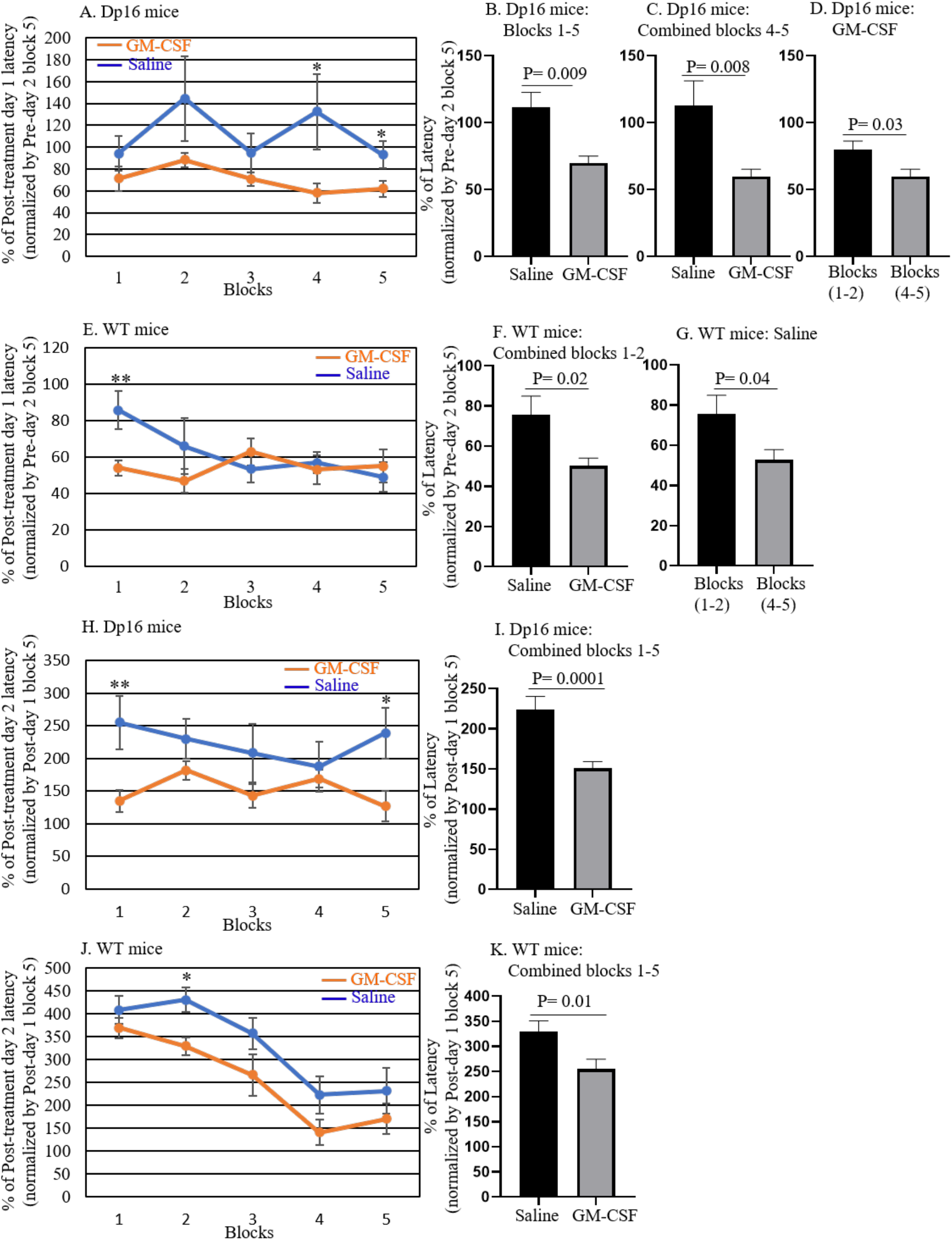
GM-CSF Treatment Improves RAWM Performance in Male Dp16 and Wild-type Mice. To evaluate the effect of GM-CSF treatment on spatial memory, latencies on post-treatment Day 1 for Dp16 mice and their WT littermates injected with saline or GM-CSF were normalized to their mean latency in block 5 of pre-treatment Day 2, and the data were then compared between saline and GM-CSF injection for each genotype. For quantitative analyses, latencies on post-treatment Day 1 were calculated relative to 100% for block 5 of pre-treatment Day 2 data. Normalized latencies were analyzed in blocks of three trials (with a total of five blocks from 15 trials) and the GM-CSF-treated group was compared with the saline-injected group of Dp16 mice (A; n=5-6) and WT mice (E; n=8-9). After averaging the normalized latencies of post-treatment Day 1 from all blocks, the mean values were compared in Dp16 mice treated with saline or with GM-CSF (B; n=5-6). Combined data from blocks 4-5 of Dp16 mice were compared for saline and GM-CSF injection (C; n=5-6). Combined data from blocks 1-2 were compared with the combined data from blocks 4-5 for Dp16 mice treated with GM-CSF (D; n=5-6). Similarly, combined data from blocks 1-2 of WT mice were compared for saline and GM-CSF injection (F; n=8-9). Combined data from blocks 1-2 were compared with the combined data from blocks 4-5 for WT mice injected with saline (G; n=8-9). The effect of GM-CSF treatment on learning flexibility was evaluated by normalizing the latency of each saline- or GM-CSF-injected mouse at post-treatment day 2 with its latency at block 5 of post-treatment day 1 and calculated data relative to 100% of block 5 post-treatment day 1 (H-K). Normalized latencies were calculated in blocks of three trials, and the GM-CSF-injected group was compared with the saline-injected group of Dp16 mice (H; n=5-6) and WT mice (J; n=8-9). Combined data from blocks 1-5 were compared for saline- and GM-CSF-injected Dp16 mice (I; n=5-6) and WT mice (K; n=8-9). Data are represented as mean ± SEM for the different groups of mice. Statistical significance was determined by the unpaired Student’s *t*-test for comparison between groups. \**p* <0.05; \*\**p* <0.01.

To assess flexibility in learning, on testing day 2 of the post-treatment RAWM task, the platform was moved (from Arm 4 to Arm 6, leaving other maze cues unchanged). We calculated the latency of each GM-CSF-treated or saline-injected mouse on post-treatment day 2 as a percentage of its latency in block 5 of post-treatment day 1. Normalized latencies were analyzed in blocks of three trials (for a total of five blocks from 15 trials), and the GM-CSF-treated group was compared with the saline-injected group. The Dp16 mice treated with GM-CSF showed significantly shorter latencies in blocks 1 and 5 (Figure 5H). Combining the normalized latencies of blocks 1-5 on post-treatment day 2 showed that the Dp16 mice treated with GM-CSF required significantly less time (shorter latencies) to reach the platform compared to the saline-injected Dp16 mice (*p*=0.0001) (Figure 5I). Similarly, GM-CSF-treated WT mice required significantly less time to reach the platform compared to saline-injected WT mice in block 2 (Figure 5J). Combining the normalized latency of blocks 1-5 in WT mice showed that, overall, GM-CSF-treated WT mice required significantly less time to reach the platform compared to saline-injected WT mice (*p*=0.01) (Figure 5K).

### GM-CSF Treatment Leads to Elevated Cytokine Levels in the Plasma of WT and Dp16 Mice

Although the effects of GM-CSF treatment on cytokine levels have been well-studied in humans and in WT mice (45, 51, 52), the effects of GM-CSF treatment on cytokine levels in Dp16 mice have not been investigated. We assessed the levels of inflammatory cytokines and metabolic proteins in the plasma of WT and Dp16 mice after 10 consecutive days of injection with GM-CSF or saline treatment using the Meso Scale Discovery (MSD) platform. We measured the levels of 15 analytes (i.e., IL-6, IL-2, TNF*α*, IP-10, MCP-1, MDC, IL-1β, IL-5, IL-22, IL-33, IFN*γ*, VEGF-A, KC/GRO, BDNF, and MMP-9) in four treatment groups: (1) WT mice injected with saline, including 3 males and 2 females; (2) WT mice injected with GM-CSF, including 4 males and 1 female; (3) Dp16 mice injected with saline, including 4 males and 1 female; and (4) Dp16 mice injected with GM-CSF, including 4 males and 1 female. Figure 6 shows the plasma levels of six key inflammation-associated cytokines (i.e., IL-6, IL-2, TNF*α*, IP-10, MCP-1, and MDC) in each of the four treatment groups. Dp16 and WT mice injected with saline did not show significant differences in the levels of these six cytokines (Figure 6A-F), but there was a trend toward increased levels of interleukin-6 (IL-6), interleukin-2 (IL-2), tumor necrosis factor *α* (TNF*α*), and macrophage-derived chemokine (MDC) in saline-injected Dp16 mice compared to saline-injected WT mice. In contrast, both WT and Dp16 mice treated with GM-CSF showed significantly higher plasma levels of IL-2 (WT mice, p=0.005; Dp16 mice, p= 0.02; Figure 6B) and MDC (WT mice, p <0.001; Dp16 mice, p= 0.03; Figure 6F) compared to saline-injected WT and Dp16 mice, respectively, whereas the plasma levels of interferon *γ*-induced protein 10 (IP-10) were significantly higher only in GM-CSF-treated Dp16 mice compared to saline-injected Dp16 mice (p=0.03; Figure 6D). Strong trends toward increased plasma levels of IL-6 (WT mice, p=0.07; Dp16 mice, p= 0.07; Figure 6A) and TNF*α* (WT mice, p=0.07; Dp16 mice, p= 0.06; Figure 6C) were observed in both WT and Dp16 mice after GM-CSF treatment. We did not observe significant differences in the plasma levels of MCP-1 (Figure 6E), or in the plasma levels of the nine other cytokines and metabolic proteins (i.e., IL-1β, IL-5, IL-22, IL-33, IFN*γ*, VEGF-A, KC/GRO, BDNF, or MMP-9; Supplementary Figure 1) in Dp16 or WT mice injected with GM-CSF or saline.

**Figure 6.**
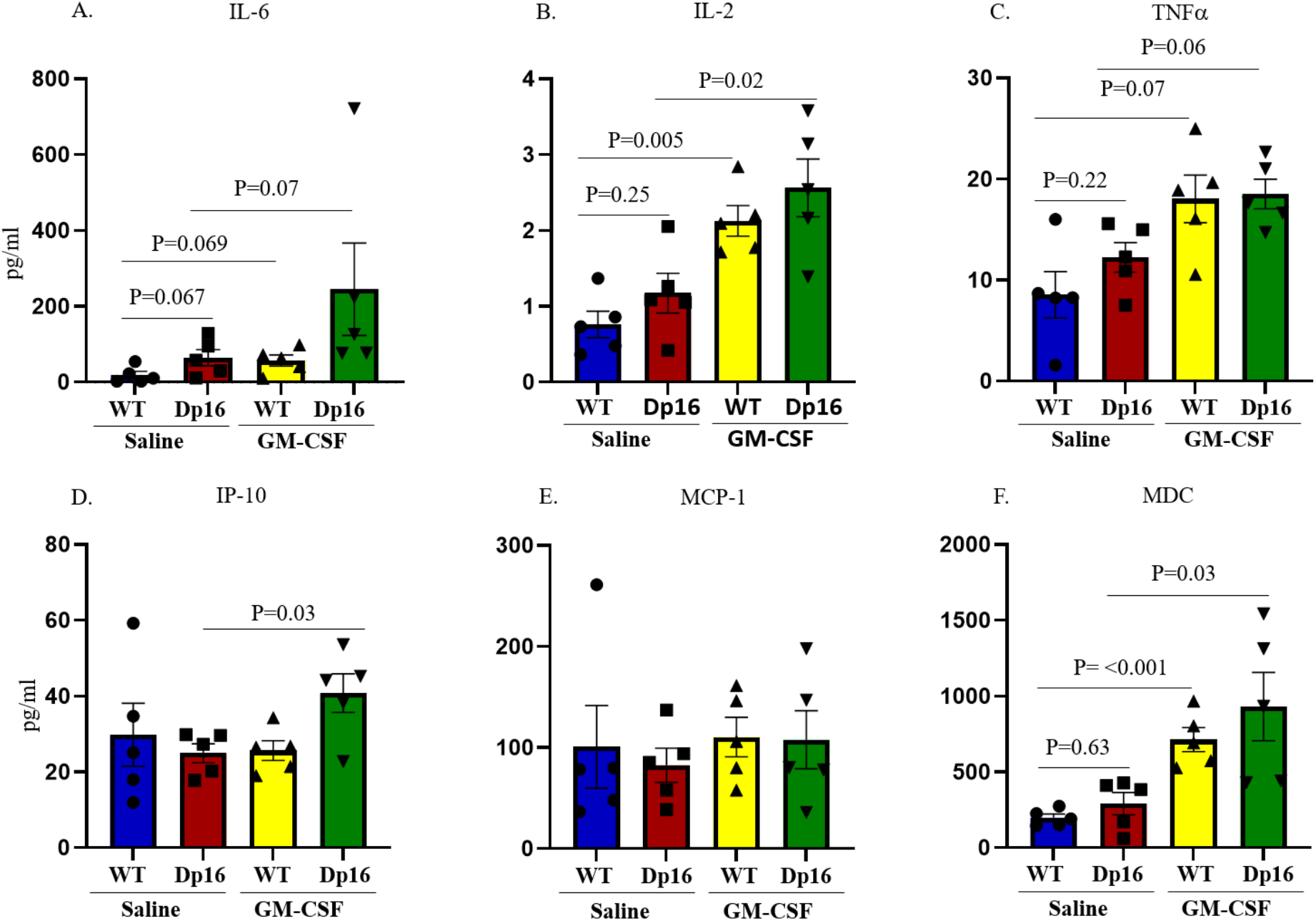
GM-CSF Treatment is Associated with Higher Plasma Levels of Cytokines in Both Wild-Type and Dp16 Mice. The Meso-Scale Discovery platform was used to determine plasma levels of six key inflammation-associated cytokines (i.e., IL-6, IL-2, TNF*α*, IP-10, MCP-1, and MDC) following GM-CSF treatment or saline injection. No significant differences were observed in the plasma levels of these six cytokines in saline-injected WT mice compared to saline-injected Dp16 mice (A-F). Treatment with GM-CSF led to significantly higher levels of interleukin-2 (IL-2) and macrophage-derived chemokine (MDC) in the plasma of both GM-CSF-treated WT and Dp16 mice compared to saline-injected WT and Dp16 mice (B & F), respectively, whereas the plasma levels of interferon *γ*-induced protein 10 (IP-10) were significantly higher only in GM-CSF-treated Dp16 mice compared to saline-injected Dp16 mice (D). In addition, there was a strong trend toward higher plasma levels of interleukin-6 (IL-6) and tumor necrosis factor *α* (TNF*α*) in the plasma of both GM-CSF-treated WT and Dp16 mice compared to saline-injected WT and Dp16 mice (A & C), respectively. No significant differences were detected in the plasma levels of MCP-1 in GM-CSF-treated or saline-injected WT or Dp16 mice (E).

### GM-CSF Treatment Partially Restores the Distribution of Hippocampal Astrocytes in Male Dp16 Mice

Glial cell abnormalities are one of the major cellular phenotypes seen in the brains of people with DS and in mouse models of DS (53, 54). We therefore investigated the expression pattern and distribution of the astrocytic marker protein, glial-fibrillary acidic protein (GFAP), in the hippocampi of male Dp16 mice and their male WT littermates. We visually counted the number of GFAP-positive clusters (blinded as to genotype and treatment). WT mice injected with saline or GM-CSF show GFAP-positive astrocytes with similar morphology and distribution (Figures 7A and 7B). In contrast, compared to WT, Dp16 mice treated with saline show GFAP-positive astrocytes with more dense cell bodies that are clustered together with crossed branches, indicating diffuse astrogliosis (Figure 7C). However, Dp16 mice treated with GM-CSF show fewer abnormal astrocyte clusters and a distribution of astrocytes with morphology more similar to that of WT (Figure 7D). This revealed significantly higher numbers of astrocyte clusters in the hippocampi of Dp16 mice injected with saline compared to WT mice (*p*=0.001). In Dp16 mice injected with GM-CSF, the astrocyte clusters were significantly reduced compared to saline-treated Dp16 mice (*p*=0.03) (Figure 7E). These results show that Dp16 mice exhibit astrocyte abnormalities in their hippocampi, and that treatment with GM-CSF partially reverses the morphological abnormalities of astrocytes and reduces astrocyte clustering in the hippocampi of Dp16 mice.

**Figure 7.**
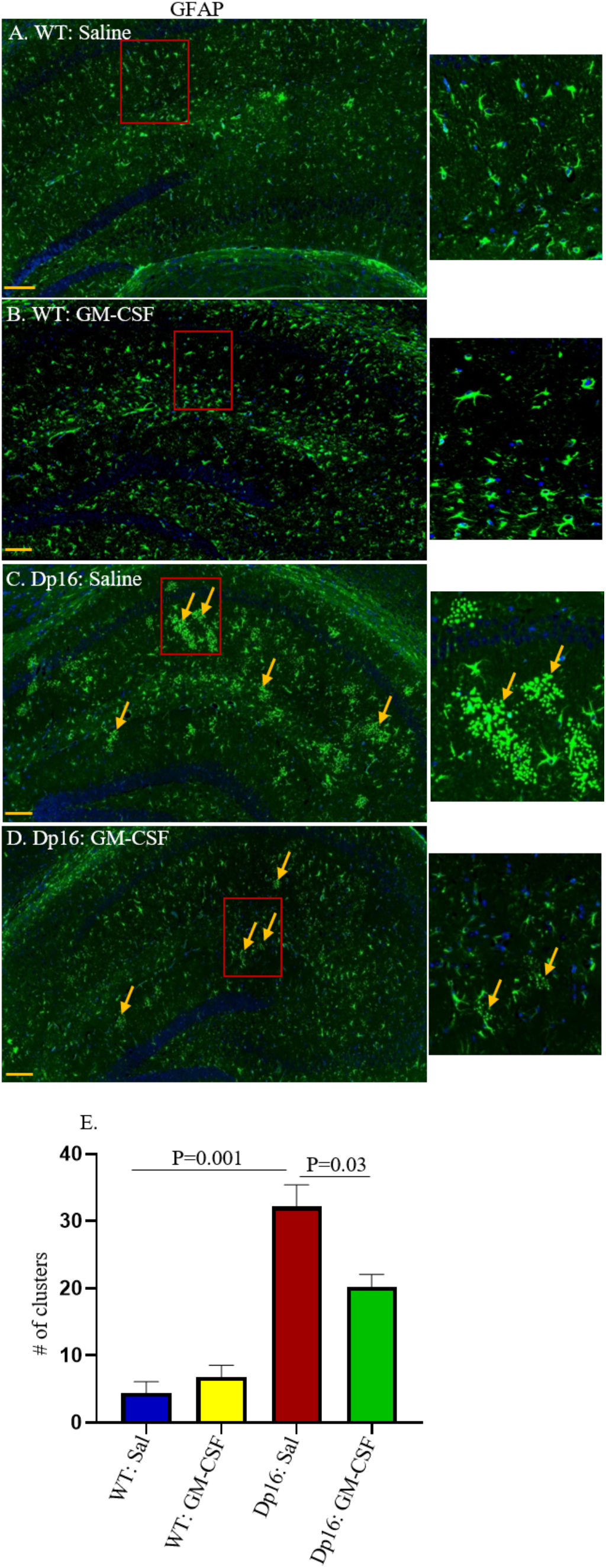
GM-CSF Treatment Partially Restores the Distribution of Hippocampal Astrocytes in Male Dp16 Mice. The patterns and distributions of glial fibrillary acidic protein (GFAP)-positive astrocytes (green) were examined in the hippocampi of male WT and Dp16 mice injected with saline or with GM-CSF. Data are shown for WT mice injected with saline (A), WT mice treated with GM-CSF (B), Dp16 mice injected with saline (C), and Dp16 mice treated with GM-CSF (D). Dp16 mice injected with saline (C) showed abnormal clusters of GFAP-positive astrocytes compared to WT mice injected with saline (A). GM-CSF treatment of WT mice had no effect on hippocampal astrocyte distribution (B) compared to WT mice injected with saline. Dp16 mice treated with GM-CSF showed significantly reduced levels of the abnormal clusters of GFAP-positive astrocytes (D) compared to Dp16 mice injected with saline (C). Arrows show abnormal clusters of GFAP-positive astrocytes. Quantitative analyses showed significantly higher numbers of abnormal clusters of GFAP-positive astrocytes in Dp16 mice injected with saline compared to WT mice injected with saline (*p*=0.001), which were reduced in Dp16 mice treated with GM-CSF (*p*=0.03) (E). For each bar, the data are represented as mean ± SEM for the separate groups of mice. Statistical significance was determined by the unpaired Student’s *t*-test for comparison between groups. Scale bar: 200 μm (20X magnification). All experiments were repeated 2-4 times with similar results (n=3 male mice in each of the four groups).

### GM-CSF Treatment Leads to an Increased Number of Calretinin-Positive Interneurons in Male Dp16 Mice

Previous studies have shown that people with DS exhibit an imbalance between excitatory and inhibitory neuronal activity in the brain, which is also seen in the Ts65Dn mouse model of DS (11, 16). Abnormalities are also observed in interneurons in induced pluripotent stem cells (iPSCs) from people with DS and from Ts65Dn mice (55, 56). We therefore investigated the numbers of calretinin-positive interneurons in the brains of male Dp16 mice and their male WT littermates injected with saline or with GM-CSF. Paraffin-embedded tissues from hippocampus and cortex were stained with an anti-calretinin antibody, and the numbers of calretinin-positive interneurons were counted. Compared to WT mice, Dp16 mice show a trend to a reduction in the number of calretinin-positive interneurons in the entorhinal cortex (*p*=0.06) (Figures 8A and 8C). GM-CSF treatment significantly increased this number (*p*=0.046) (Figures 8D and 8E), to levels indistinguishable from those of their saline-treated WT littermates. Similarly, the number of calretinin-positive interneurons was significantly reduced (*p*=0.01) in the subiculum of Dp16 mice injected with saline compared to WT mice (Figures 8F, 8H, and 8I). This number was significantly increased in Dp16 mice injected with GM-CSF (*p*=0.04) (Figure 8I). No significant differences were observed between WT mice treated with saline or with GM-CSF in either the entorhinal cortex or the subiculum. Together, these results show that GM-CSF treatment is able to partially reverse the deficit in the number of interneurons in Dp16 male mice.

**Figure 8.**
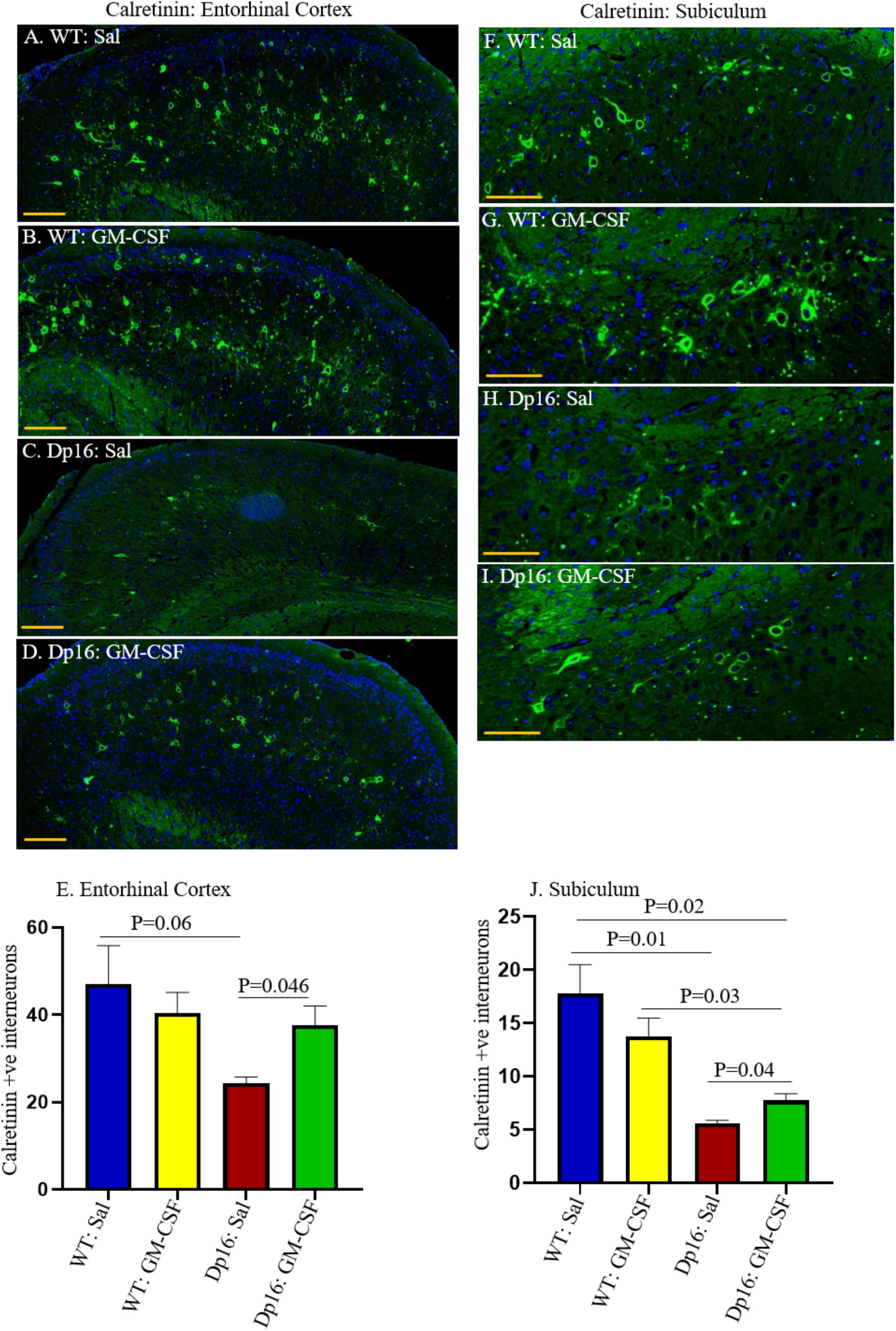
GM-CSF Treatment Partially Restores the Number of Calretinin-Positive Interneurons in Male Dp16 Mice. The patterns and distributions of calretinin-positive interneurons (green) were examined in the entorhinal cortex (EC) of male WT mice injected with saline (A), male WT mice treated with GM-CSF (B), male Dp16 mice injected with saline (C), and male Dp16 mice treated with GM-CSF (D). Quantitative analyses showed a trend towards a lower number of calretinin-positive interneurons in the entorhinal cortex of Dp16 mice injected with saline compared to WT mice injected with saline (*p*=0.06).The number of calretinin-positive interneurons in the entorhinal cortex of the Dp16 mice was significantly increased following GM-CSF treatment compared to Dp16 mice injected with saline (*p*=0.0406), and was statistically no different than in WT mice injected with either saline or GM-CSF (E). The expression patterns and distributions of calretinin-positive interneurons (green) were also examined in the subiculum of male WT mice injected with saline (F), male WT mice treated with GM-CSF (G), male Dp16 mice injected with saline (H), and male Dp16 mice treated with GM-CSF (I). There was a significantly reduced number of calretinin-positive interneurons in the subiculum of Dp16 mice injected with saline compared to WT mice (*p*=0.01). The number of calretinin-positive interneurons in the subiculum of Dp16 mice was significantly increased following GM-CSF treatment (*p*=0.04), but it was still lower than in WT mice injected with saline or with GM-CSF (J). For each bar, data are represented as mean ± SEM for separate groups of mice. Statistical significance was determined by the unpaired Student’s *t*-test for comparison between groups. Scale bar: A-D=100 μm (20X magnification) and F-I=50 μm (20X magnification). All experiments were repeated 2-4 times with similar results (n=3 male mice in each of the four groups).

## Discussion

We analyzed 12-14 month-old male and female Dp16 mice and their age- and sex-matched littermates in the Open Field and alternating Y-maze tasks at baseline, followed by an investigation of the effects of treatment with GM-CSF, which is an innate immune system stimulator and pro-inflammatory cytokine, on learning/memory in the RAWM and on brain pathology in male mice. Examining both sexes, when possible, is ideal because sex differences have been noted in the descriptions of cognitive phenotypes observed in people with DS (57, 58), and sex differences are also frequently observed with drug responses in the typical population (59). However, sex differences have not been commonly investigated in DS mouse models, due in part to the fact that the popular Ts65Dn mouse model of DS breeds poorly, and that the male Ts65Dn mice are largely sterile, so that female Ts65Dn mice are generally saved as breeders. However, where investigated, sex differences are evident at the behavioral level (36). Our data show that both male and female Dp16 mice were hyperactive in the Open Field task at baseline, confirming a previous report of increased locomotor activity in male Dp16 mice of a similar age (60). However, this may be an age-dependent phenotype, because younger (2-3 month-old) male Dp16 mice did not show hyperactivity (61). Similarly, increased locomotor activity in the Open Field arena was reported in 9-12 month-old Ts65Dn mice (50), and human studies have shown that children with DS are hyperactive compared to typical children of a similar age (62, 63). We also observed sex differences in the Y-maze: at baseline, female Dp16 mice were impaired, performing no better than chance, whereas male Dp16 mice were not impaired.

Only male mice were analyzed in the RAWM, because the female Dp16 mice at this age are poor swimmers. Male Dp16 mice learned more slowly compared to their male WT littermates, indicating impaired hippocampal function. This finding is consistent with previous reports of impairment of Dp16 mice in other tasks that require hippocampal function, e.g., Morris Water Maze and Contextual Fear Conditioning, and of hippocampal learning deficits in other mouse models of DS, including Ts65Dn and Ts1Cje mice, each of which is trisomic for a subset of the genes that are trisomic in the Dp16 mice (56, 61, 64–67). This finding is also consistent with previous studies that showed that Dp16 mice display abnormal hippocampus-based long-term potentiation (LTP) (60, 61, 67, 68).

We found that treatment with GM-CSF increased the levels of some pro-inflammatory cytokines in the plasma of WT and Dp16 mice, supporting its inflammatory role in both groups of mice. In contrast to the commonly held hypotheses that blocking or reversing the activated innate immune system and chronic inflammation associated with DS may reverse many of its features, our results show that treatment with the pro-inflammatory cytokine GM-CSF effectively ameliorated both learning deficits and brain pathology in the Dp16 mouse model of DS. Specifically, daily subcutaneous injection of GM-CSF rescued male Dp16 deficits in the RAWM, improving their performance to a level comparable to that of age-matched, saline-treated male WT littermates. GM-CSF treatment also enhanced RAWM performance in middle-aged WT littermates, as we and others have shown previously (41, 43). Additionally, when the platform was moved to a new arm on testing day 2 of the post-treatment RAWM task, we found that GM-CSF treatment improved learning flexibility in both WT and Dp16 mice. Impaired cognitive flexibility is often observed among individuals with DS (69). Although studies have shown that GM-CSF treatment improves learning/memory and removes amyloid plaques in mouse models of AD, Dp16 mice do not develop AD amyloid pathology at any age. Therefore, GM-CSF treatment must lead to improved learning/memory in Dp16 mice and WT mice via an amyloid-independent mechanism(s), possibly related to its pro-inflammatory activity. It is perhaps relevant that GM-CSF has been shown to increase the number of T-regulatory cells (Tregs) in mouse models of Parkinson’s disease (PD), in mouse models of other autoimmune and inflammatory conditions, and in participants in a human PD clinical trial (70–76). Treg levels are deficient in people with DS, and people with DS are susceptible to autoimmune diseases (77, 78), which has been attributed to defective Tregs (78). For example, autoimmune thyroidism or Hashimoto’s thyroiditis (HT) is prevalent in people with DS (79, 80), and, indeed, GM-CSF-stimulated Treg induction has been shown in several studies to inhibit experimental autoimmune thyroiditis, a common model of HT (75, 77, 81, 82).

### Effects of GM-CSF on Neuronal Function

Despite its known pro-inflammatory properties, GM-CSF also affects multiple CNS processes that are consistent with and may shed light on its unexpected beneficial effects on learning/memory in the Dp16 mouse model of DS. For example, we found that the density of cells that stain positively for calretinin, a marker for interneurons, is decreased in the entorhinal cortex and in the subiculum of male Dp16 mice, and that GM-CSF treatment leads to a partial restoration of these numbers. Reduced density of calretinin-positive interneurons has also been seen in induced pluripotent stem cells (iPSCs) from people with DS (55). Because inhibitory interneurons contribute to neuronal activity, information processing for neuronal networks, synaptic plasticity, and learning/memory (83–85), their impaired function might impact cognitive processing in the brains of people with DS. However, reduced interneuron numbers cannot fully explain the learning/memory deficits in Dp16 mice because, in contrast to our observations in 12-14 month-old mice, in 4 month-old Dp16 mice, the density of calretinin-positive interneurons was normal in the striatum oriens of the hippocampus (61, 66). That 2-4 month-old Dp16 mice show impairment in other hippocampal-based tasks but show no defects in the RAWM suggests that other cellular abnormalities underlie some of the learning/memory deficits at this age. Furthermore, because we did not find that GM-CSF increases interneuron density in 12-14 month-old WT mice, the improved learning/memory function observed in WT mice must occur via a different mechanism(s).

Other studies support a beneficial effect of GM-CSF on recovery from neuronal damage or dysfunction, modulation of apoptosis-related genes and prevention of programmed neuronal cell death, restoration of cerebral blood flow, and promotion of axonal growth, regeneration, and survival in animal models of several CNS-related disorders (86–94). (95–97).

Additionally, GM-CSF is a growth factor for neural stem cells (NSCs) and prevents neuronal apoptosis (98–103), with effects only on cell survival at low concentrations and with effects on cell survival, proliferation, differentiation, and functional activation at higher concentrations (104). GM-CSF can cross the blood-brain barrier (105), and is also produced within the brain, where numerous cell types express the GM-CSF receptor, including neurons, oligodendrocytes, microglia, astroglia, and endothelial cells (106, 107). Indeed, GM-CSF has also been shown to play a major role in neuronal plasticity that is critical for learning/memory (108). In addition to its effects on NSCs, GM-CSF has been shown to mobilize mesenchymal stem cells (MSCs) from the bone marrow and to enhance their differentiation and migration in response to injury (109, 110). MSCs have been reported to have beneficial cognitive and regenerative effects and are being investigated as potential positive effectors in several neurological disorders, including AD, PD, and spinal cord injury (111–117).

We previously showed neurological benefits of GM-CSF treatment in a mouse model of AD that was also unexpected, because the well-documented existence of inflammation and its contribution to AD led many to predict that, in general, non-steroidal anti-inflammatory drugs (NSAIDs) would be protective against AD, and that a pro-inflammatory molecule such as GM-CSF would be expected to exacerbate AD (118–122). In fact, the NSAID clinical trials in AD and mild cognitive impairment (MCI) participants failed to show any reversal or slowing of disease symptoms and may have led to accelerated cognitive decline (123, 124). Furthermore, treatment of a mouse model of AD with GM-CSF reversed cognitive deficits in the RAWM, reduced amyloidosis, and increased synaptophysin expression and microglial density (41). Similar to the current study, cognitive improvement was also reported in age-matched WT mice treated with GM-CSF (41). These results showing improved cognition in WT and AD mice treated with GM-CSF and reversal of neuropathology in AD mice have been replicated by other groups (42, 43). Our recently completed clinical trial in which mild-to-moderate AD participants were treated with recombinant human GM-CSF (sargramostim/Leukine^®^) for 15 days showed that sargramostim treatment was safe, led to significantly improved MMSE scores, and partly restored levels of plasma biomarkers of AD brain pathology and neuronal damage (e.g., Aβ40 and Aβ42, total tau, and ubiquitin C-terminal hydrolase L1 [UCH-L1]) towards normal. People with DS before the onset of AD pathology also exhibit increased brain levels of Tau and low levels of UCH-L1(125). We are collaborating to measure these proteins in the plasma of people with DS throughout the lifespan, including at ages before AD-related brain pathology is present, compared to typical age-matched controls.

### Role of Astrocytes in Learning/Memory

Astrocytes play an essential supportive role with regard to neuronal function, both directly and indirectly, by regulating glucose metabolism, controlling cerebral blood flow, and maintaining homeostasis of ion and neurotransmitter levels in the CNS (reviewed in (126)), as well as by playing pivotal roles in information processing, brain wiring, and cellular programming (127–129). Astrocytes modulate synaptic activity, regulate synapse formation, transmission, and plasticity, and are required for the maintenance of LTP via their interactions with neurons and/or with other astrocytes (130). Aberrations in the generation, number, morphology, and proliferation of astrocytes are observed in many neurodegenerative and neuropsychiatric disorders, including AD, schizophrenia, autism, Rett’s syndrome, and epileptic and non-epileptic seizures (131–133). Numerous studies have also described abnormalities in the numbers and morphologies of astrocytes in people with DS (12, 13, 53, 134, 135). Our results showed that GFAP-expressing astrocytes in the hippocampi of male Dp16 mice are morphologically abnormal and that GM-CSF treatment partially reversed these abnormalities, correlating with improvements in the RAWM task. In agreement with this finding, studies of spinal cord injury showed that GM-CSF treatment reduced GFAP astrogliosis, inhibited glial scar formation, prevented neuronal cell death, preserved axon and myelin structure and facilitated axonal regeneration, and promoted functional recovery (136, 137). WT littermates in our studies also showed improved RAWM performance with GM-CSF treatment without changes in GFAP staining, suggesting that the GM-CSF-induced improvements occur via alternative mechanisms.

In summary, these data show that treatment with the pro-inflammatory cytokine GM-CSF improves learning/memory and neuropathology in the Dp16 mouse model and improves learning/memory in WT mice. However, as discussed, multiple studies have shown that people with DS and also aged people (138–141) exhibit an auto-inflammatory or ‘inflammaging’ syndrome that would seem to preclude any benefit from GM-CSF treatment. However, it is more likely that the dichotomy may be more apparent than real because GM-CSF is not merely a pro-inflammatory molecule. A more accurate description would be that GM-CSF modulates the innate immune system, especially in the setting of immune system dysregulation and in the brain (41, 77, 142, 143). Indeed, GM-CSF treatment not only increases the levels of many cellular and cytokine biomarkers of inflammation in the blood of mild-to-moderate AD patients (e.g., neutrophils, monocytes, lymphocytes, IL-2, IL-6, and TNF*α*), but also reduces the level of the inflammatory cytokine IL-8 and increases the level of the typically anti-inflammatory cytokine IL-10 (45). Thus, GM-CSF has a much more complex physiological effect than simply being pro-inflammatory. Furthermore, suppressing the inflammation associated with DS in the periphery may be beneficial in the setting of certain acute disorders (40). Thus, the results reported here, although somewhat unexpected, contribute to our growing understanding of the complexity of the innate immune system in the context of inflammation in people with DS and in typical aging populations.

## Conclusions

The population of people with DS continues to expand, and the average lifespan of people with DS continues to increase. There is, therefore, a critical need to develop therapeutics to improve cognitive function in people with DS and to help them to live more independently. In contrast to previous expectations that inhibiting inflammation and the innate immune system would be the most effective therapy for co-morbidities of DS, we report that treatment with GM-CSF, often described as pro-inflammatory, which we have also confirmed in the Dp16 mice, reverses learning/memory deficits in a hippocampal-based task, ameliorates astrocyte abnormalities, and partially normalizes the levels of interneurons in 12-14 month-old male Dp16 mice, while also improving cognitive function in their WT littermates. These findings were unexpected in view of the extensive body of literature describing abnormal and chronic inflammatory perturbations across the lifespan in people with DS and in the typical aging population. Collectively, our results should open the possibility that the field’s current approach to cognitive therapy in people with DS, as well as in the typical aging population, should be reconsidered, reversed, or at least modified. Furthermore, these new results, taken together with previous findings of the benefits of GM-CSF treatment in AD mice and other models of neurological disease, and of our results showing GM-CSF/sargramostim efficacy with regard to cognition in clinical trial participants with mild-to-moderate AD as well as in cancer patients, combine to support the hypothesis that GM-CSF may promote neuronal recovery from injury or neurological disease through multiple mechanisms, some of which may also enhance cognitive function. The beneficial effects of GM-CSF on learning and memory may reflect its pro-inflammatory activity, its other physiological effects, or both. These results indicate that with an almost 30-year history of safety in numerous patient populations across the lifespan, including in vulnerable populations, sargramostim should be tested for safety, tolerability, and efficacy for improving cognition in people with DS.

## Acknowledgements

Research support was provided by NIH grant R01 AG037942-01A1, the State of Colorado, the University of Colorado School of Medicine, the Linda Crnic Institute for Down Syndrome, Don and Sue Fisher, the Hewit Family Foundation, Marcy and Bruce Benson, the Global Down Syndrome Foundation, and other generous philanthropists. In addition, we would like to thank the Animal Behavior Core, which is supported by NIH grant 5P30NS048154, and Kim Jordan, Ph.D. and the Human Immunology & Immunotherapy Initiative at the University of Colorado Anschutz Medical Campus

## Author Contributions

Conceptualization, T.D.B. and H.P.; Methodology, M.M.A. T.D.B., and H.P.; Investigation, M.M.A., A.CJ.W., M.E., D.A.S., C.C., L.A., P.A., N.M., and V.A.; Formal Analysis, M.M.A., S.S., and H.P.; Writing – Original Draft, M.M.A. and T.D.B.; Writing – Review & Editing, M.M.A., T.D.B., H.J.C., K.J.G., and H.P.; Resources, K.J.G., and H.P.; Visualization, M.M.A.; Project Administration, T.D.B.; Supervision, H.P.; Funding Acquisition, H.P.

---

**Supplementary Figure 1.**
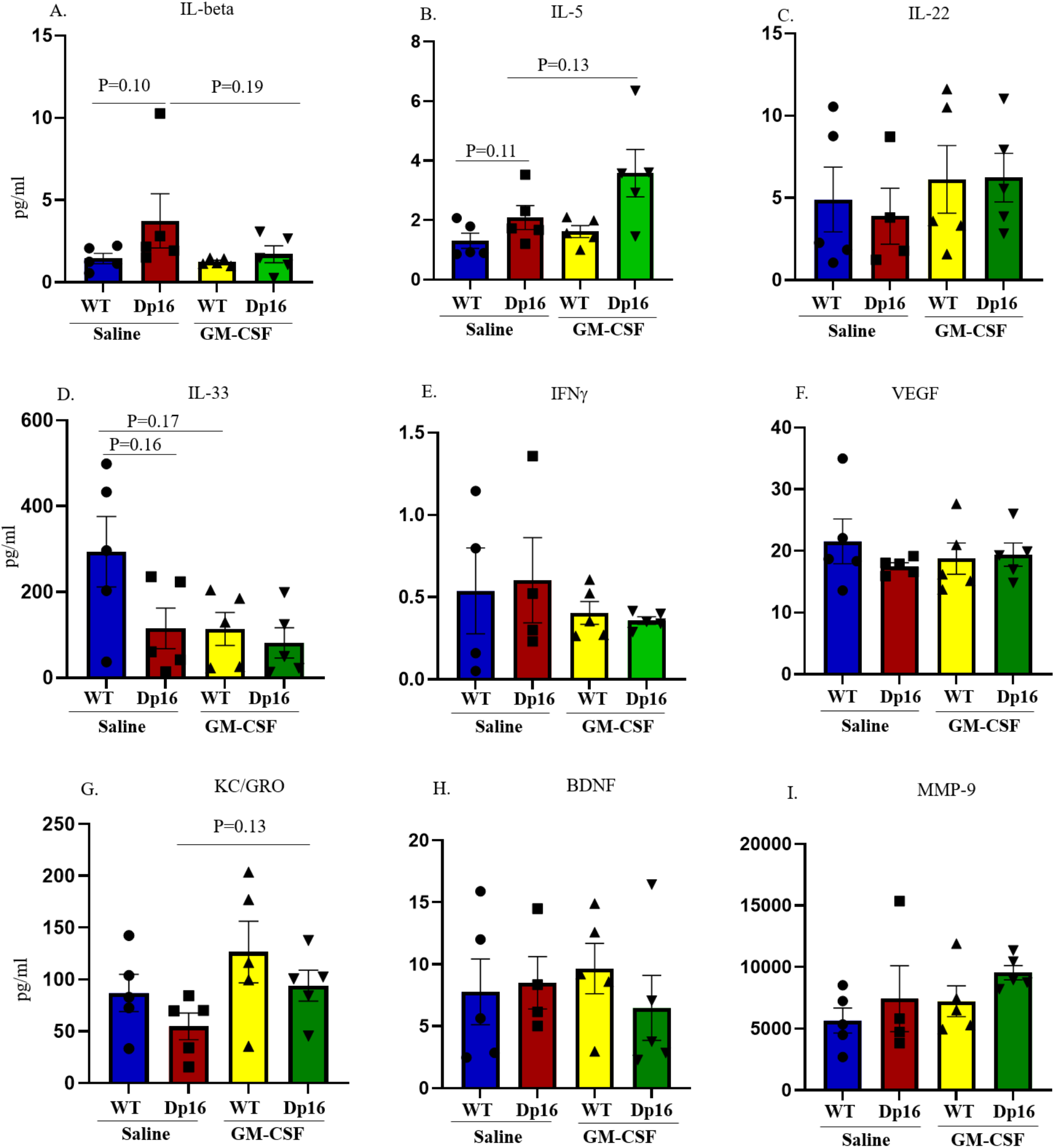
Plasma Levels of Additional Cytokines and Metabolic Proteins in WT and Dp16 mice After GM-CSF Treatment or Saline Injection The Meso-Scale Discovery (MSD) platform was used to determine the levels of additional cytokines and metabolic proteins in the plasma of both WT and Dp16 mice after GM-CSF treatment or saline injection (A-I). No significant differences were observed in the levels of IL-beta, IL-5, IL-22, IL-33, IFN*γ*, VEGF-A, KC/GRO, BDNF, or MMP-9 after either GM-CSF treatment or saline injection in WT or Dp16 mice.

